# MCT2 Overexpression Rescues Metabolic Vulnerability and Protects Retinal Ganglion Cells in Two Models of Glaucoma

**DOI:** 10.1101/2020.02.14.950097

**Authors:** Mohammad Harun-Or-Rashid, Nathaniel Pappenhagen, Ryan Zubricky, Lucy Coughlin, Assraa Hassan Jassim, Denise M. Inman

## Abstract

Improving cellular access to energy substrates is one strategy to overcome observed declines in energy production and utilization in the aged and pathologic central nervous system. Monocarboxylate transporters (MCTs), the movers of lactate, pyruvate, and ketone bodies into or out of a cell, are significantly decreased in the DBA/2J mouse model of glaucoma. In order to confirm MCT decreases are disease-associated, we decreased MCT2 in the retinas of MCT2^fl/+^ mice using an injection of AAV2-cre, observing significant decline in ATP production and visual evoked potential. Restoring MCT2 levels in retinal ganglion cells (RGCs) via intraocular injection of AAV2-GFP-MCT2 in two models of glaucoma, the DBA/2J (D2), and a magnetic bead model of ocular hypertension (OHT), preserved RGCs and their function. Viral-mediated overexpression of MCT2 increased RGC density and axon number, reduced energy imbalance, and increased mitochondrial function as measured by cytochrome c oxidase and succinate dehydrogenase activity in both models of glaucoma. Ocular hypertensive mice injected with AAV2:MCT2 had significantly greater P1 amplitude as measured by pattern electroretinogram than mice with OHT alone. These findings indicate overexpression of MCT2 improves energy homeostasis in the glaucomatous visual system, suggesting that expanding energy input options for cells is a viable option to combat neurodegeneration.

**Highlights:**

- Loss of MCT2 in retina compromises visual function and ATP production in optic nerve
- MCT2 overexpression preserves RGC soma and axon number in two glaucoma models
- MCT2 overexpression improves energy homeostasis in optic nerve

## 1. Introduction

Glaucoma is an optic nerve degeneration accompanied by vision loss believed to be a result of a progressive loss of retinal ganglion cell axons followed by their cell bodies (Buckingham et al., 2008; Howell et al., 2008). Indeed, it is this window between axonopathy and soma loss that may present a therapeutic window for glaucoma treatment (Crish et al., 2010). In the DBA/2J (D2) mouse model of glaucoma, optic nerves (ONs) are predisposed to neuropathy through metabolic deficiencies revealed through decreased resilience to oxygen-glucose deprivation (Baltan et al., 2010) and limits on metabolic switching when challenged (Jassim et al., 2019). Metabolic deficiencies in particular have shown promise as a potential target in the effort to reverse glaucoma (Harun-or-Rashid et al., 2018; Williams et al., 2017).

Glucose is the primary energy source for the mammalian brain (Siesjö, 1978), yet neurons can utilize lactate for oxidative metabolism. Astrocytes produce lactate as a glycolytic end-product which can be transported to neurons for use (Bouzier-Sore et al., 2006) via the monocarboxylate transporter (MCT) family (Halestrap, 2013a; Halestrap and Meredith, 2004; Simpson et al., 2007). A gradient of lactate in brain parenchyma enables lactate flux from astrocytes to neurons (Mächler et al., 2016). In locations such as the optic nerve, axons are provided this lactate from oligodendrocytes (Lee et al., 2012; Saab et al., 2016). Given the positioning of astrocytes between CNS vasculature and the nodes of Ranvier of ON axons (Dutta et al., 2018; Serwanski et al., 2017), it is possible that astrocytes are privy to axon energy needs and can supply glucose or lactate as well.

Within the MCT family, only MCT1 through 4 have been shown to transport monocarboxylates relevant to metabolism (Halestrap and Meredith, 2004). MCT1 is widely expressed early in postnatal development, and expression generally increases with age in astrocytes (Simpson et al., 2007). MCT2 is localized to neurons in the adult mouse brain and associated with the postsynaptic density protein PSD-95 (Pierre et al., 2002; Rafiki et al., 2003; Simpson et al., 2007). In the rat retina, MCT2 is heavily expressed throughout the inner retina, including the inner plexiform layer (Gerhart et al., 1999). Ancillary proteins such as basigin, embigin, and the neuroplastins Np55 and Np65 are necessary for MCT-1, −2, and −4 localization, chaperoning the transporters to the cell surface (Halestrap, 2013a; Wilson et al., 2013). MCT2 preferentially binds embigin (Halestrap, 2013b; Wilson et al., 2005). The chaperone proteins do not affect MCT lactate affinity (Ovens et al., 2010), but their association with the MCTs is necessary for maintaining activity (Manoharan et al., 2006). Of these four monocarboxylate transporters, our interest primarily lies with MCT2 because it is the main transporter for monocarboxylates to neurons. Inhibition of MCTs results in decreased retinal function with reductions in phototransduction sensitivity and reduction in neuronal levels of the neurotransmitter glutamate (B. V Bui et al., 2004). However, inhibition of MCTs caused no changes in retinal function when preceded by ischemia (Melena et al., 2003). Ischemia/hypoxia leads to increases in glucose transporters (Yang et al., 2014), possibly decreasing the need for lactate transport, at least for fuel. Both MCT1 and MCT2 protein were significantly decreased in the D2 optic nerve despite stable axon number and glial hypertrophy (Harun-or-Rashid et al., 2018). It is thus hypothesized that MCT2 may be a therapeutic target where rescue of glaucomatous axons and protection of vision may be possible.

Monocarboxylate transporters, including MCT1 and MCT2, are upregulated in the D2 optic nerve after two months of the ketogenic diet (Harun-or-Rashid et al., 2018). A ketogenic diet is a low-carbohydrate, high-fat, and calorie-restricted diet used to treat childhood epilepsies that are refractory to medication therapy. The ketogenic diet upregulated oxidative phosphorylation in mice, increased mitochondrial proteins and mitochondrial biogenesis, and elevated the phosphocreatine/creatine ratio (Bough et al., 2006; Harun-or-Rashid et al., 2018). This is consistent with studies that have shown that increases in lactate and the ketone body β-hydroxybutyrate are associated with increases in levels of MCT2 protein and mRNA (Takimoto and Hamada, 2014). This upregulation occurs at the level of translation through the stimulation of the PI3-kinase–AKT–mTOR –p70S6K pathway (Chenal et al., 2007; Harun-or-Rashid et al., 2018). MCT2 is the primary transporter for ketone bodies as well as lactate to neurons (Pierre et al., 2002; Simpson et al., 2007). As a result, it follows that the energy management of the neurons would be improved in glaucomatous conditions if more MCT2 were available. The question arises of whether the ketogenic diet is necessary or if upregulation of MCTs alone is sufficient to provide this protective effect for glaucoma.

We utilized adeno-associated virus (AAV-2) infection to increase MCT2 protein in two models of glaucoma, testing whether increasing MCT2 protein, and potentially monocarboxylate transport, could rescue metabolically challenged retinal ganglion cells (RGCs). We observed significant protection of RGCs and their axons, improved energy balance, mitochondrial function, and visual function after AAV2-directed MCT2 upregulation.

## 2. Materials and Methods

### 2.1 Animals

DBA/2J (D2), DBA/2J-Gpnmb+ (D2G), Tg(Thy1-cre/ERT2,-EYFP)HGfng/PyngJ (Thy1-cre) and B6.Cg-Gt(ROSA)26Sortm9(CAG-tdTomato)Hze/J (Rosa) mice were purchased from Jackson Laboratories (Bar Harbor, ME). MCT2^fl/fl^ mice were a generous gift from Dr. David Gozal at the University of Chicago. All mice were bred and housed at Northeast Ohio Medical University. Both sexes of mice were used. The iris stroma, due to mutations in the *Tyrp1* and *Gpnmb* genes, atrophies in the D2 mouse, leading to blockage of the aqueous humor outflow pathways and development of glaucoma. In the D2G mouse, the wildtype *Gpnmb* allele ensures the pigment dispersion does not occur, so glaucoma does not develop (Howell et al., 2007). We crossed Thy1-cre and Rosa mice to generate progeny expressing tdTomato, a red fluorescent protein, in Thy1-positive cells, using these mice for our model of ocular hypertension. All procedures were performed in accordance with the National Institutes of Health guide for the care and use of Laboratory animals (NIH Publications No. 8023, revised 1978), with approval from the Institutional Animal Use and Care Committee.

### 2.2 Intraocular Pressure

A TonoLab rebound tonometer (Tiolat-Oy, Finland) calibrated for mice was used to measure intraocular pressure (IOP). Mice were lightly anesthetized with 2.5% isoflurane, then 10-20 measures from each eye were taken within 3 minutes to avoid any anesthetic-associated reduction of IOP. D2 mice had IOP measured at 10 months of age, then again at 12 months, prior to sacrifice. IOP measurements were taken prior to ocular hypertension (OHT) to establish a baseline IOP, then IOP was measured weekly after OHT. For IOP integral calculation, the difference between the baseline and weekly IOP was multiplied by the days between measurements then summed.

### 2.3 Diets

All mice, with the exception of a subset of 9-month-old D2 mice, were fed standard rodent chow (Formulab Diet 5008) *ad libitum*. The subset of D2 mice were fed a very low carbohydrate ketogenic diet (D12369B, Research Diets, New Brunswick, NJ) for 8 weeks. The standard diet was 26.8% protein, 56.4% carbohydrate, and 16.7% fat, while the ketogenic diet was 10.4% protein, 0.1% carbohydrate, and 89.5% fat. Mice undergoing diet transition were weighed prior to changing diets, then weekly while on the ketogenic diet. Food weight was also recorded to calculate consumption. Weights and food intake for mice on the ketogenic diet are in Table 1.

**Table 1.** Indices for DBA/2J Mice on the Ketogenic or Control Diets.

| | Baseline IOP (mmHg) | Terminal IOP (mmHg) | Baseline Weight (g) | Terminal Weight (g) | Food Intake by Week (g) | kCal by Week | Plasma β-HB (mM) | Blood Sugar (mg/dL) |
| --- | --- | --- | --- | --- | --- | --- | --- | --- |
| AAV2:MCT2 D2 Keto | 16.8 ± 0.8 | 21.2 ± 0.9 | 29.7 ± 1.0 | 36.8 ± 1.4 | 12.9 ± 0.8 | 87.4 ± 5.9 | 0.753 ± 0.104 | 113.6 ± 6.6 |
| AAV2:MCT2 D2 Control | 17.1 ± 0.6 | 21.0 ± 0.7 | 30.2 ± 0.8 | 31.1 ± 0.8 | 20.1 ± 0.5 | 78.8 ± 2.0 | 0.062 ± 0.017 | 103.7 ± 6.3 |

± SEM

### 2.4 Virus

An AAV2.cre.GFP (1.0×10^13^ GC/mL; Vector Biolabs, Malvern, PA) was injected into MCT2^fl/fl^ mouse eyes to remove the flox’d MCT2 gene through cre-mediated recombination, thereby eliminating MCT2 in the infected retinal cells. The AAV2 vector carrying MCT2, AAV2.CAG2.mSLC16A7-2A.eGFP, was also produced by Vector Biolabs. AAV2:MCT2 titer was 3.1×10^12^ GC/mL. The control virus, AAV2.CMV.PI.eGFP.WPRE.bGH, was produced by the University of Pennsylvania Penn Vector Core. AAV2:eGFP titer was 4.4×10^12^ GC/mL. Each injected eye received 2µL of virus.

### 2.5 Fundus Photography

The Micron IV (Phoenix Technology Group) system was used to image the fundus in experimental and control mice after AAV2 injection. Mice were anesthetized with a Ketamine (100mg/kg)-Xylazine (10mg/kg) cocktail, then given 0.5% Tropicamide (Henry Schein Animal Health) eyedrops to dilate the pupils. GenTeal eye lubricant (Alcon) was placed on each eye for imaging. Each viral vector carried an enhanced GFP reporter, so by imaging at 400nm excitation and 509nm emission, we were able to visualize AAV2-infected cells in the ganglion cell layer.

### 2.6 Vision Testing

The Diagnosys visual testing system (Celeris) was used to measure visual evoked potential and pattern electroretinogram. For both, mice were dark adapted for ≥1h, then anesthetized with Ketamine (100mg/kg)-Xylazine (10mg/kg) and placed prone on a heated stage. For visual evoked potential (VEP), Electrodes were placed in the mouse’s cheek (reference), scalp at the base of the skull, and the base of the tail (ground). Stimulators were placed on each eye and the output of 600 sweeps was processed to yield a waveform with two positive peaks (P1 and P2) and one negative peak (N1), from which a response amplitude was calculated (Absolute value of N1 + P2). For pattern electroretinogram (PERG), an integrated electrode and stimulator with moving bar was placed on either eye while the contralateral eye held an electrode measuring background activity. One hundred sweeps yielded waveforms with a negative peak (N1), a positive peak (P1) and a second negative (N2). We report P1 and N2 amplitude in µV.

### 2.7 Seahorse Analyzer

MCT2^fl/+^ mice injected with AAV2-cre had their optic nerves removed and prepped for analysis in the Seahorse XF24 Analyzer as described (Jassim et al., 2019). The optic nerves were secured in an Islet capture microplate, within a plasma-thrombin clot, with DMEM (10mM glucose, 0.5mM sodium pyruvate, 4mM glutamine) as media. Optic nerves were subjected to the mitochondrial stress test (sequential addition of 10µg/mL oligomycin, 4µM FCCP, then 10µM antimycin-A), with either glucose or 2-deoxyglucose supplementation prior to oligomycin addition. The Analyzer measured oxygen consumption rate and extracellular acidification rate for each optic nerve.

### 2.8 Ocular Hypertension

Mice were anesthetized with isoflurane (2.5%), and 0.5% Tropicamide was placed on each cornea to dilate the pupils. Proparacaine hydrochloride (0.5%, Henry Schein Animal Health) was placed on the cornea after pupil dilation as a local anesthetic. Curved forceps were used to proptose and stabilize the eye while a glass pulled micropipette was used to inject 1.5µL of 8µL magnetic microbeads (UMC4F, Bangs Laboratories) into the central anterior chamber. Before the micropipette was removed, a neodymium magnet was used to pull the beads into the iridocorneal angle to occlude the trabecular meshwork. The micropipette was removed after the beads were distributed, and the mice were allowed to recover on a warm water blanket with Pura-Lube ophthalmic ointment on their eyes. There was no further use of the magnet after the initial injection. Two days after injection, IOP was measured as above and weekly thereafter for 4-5 weeks. Once the IOP integral reached 200mmHg-days, the mice were sacrificed, and tissue was analyzed.

### 2.9 Optic Nerve Analysis

Mice were given an overdose of sodium pentobarbital (300mg/kg, Beuthanasia-D). Optic nerves (ON) were removed and fixed by immersion in 2%paraforaldehyde with 2.5% glutaraldehyde in 0.1M sodium cacodylate buffer (pH 7.4). ON were stained with 2% osmium (Electron Microscopy Sciences), rinsed in sodium cacodylate, then taken through a series of graded ethanols to a 1:1 mixture of ethanol and propylene oxide, and finally, increasing concentrations of Ardalite 502/PolyBed^®^ (02595-1, Polysciences, Inc) in propylene oxide. Nerves were incubated under vacuum in Ardalite 502/PolyBed^®^ with three changes of embedding medium, until they were placed in a final embedding mold and cured in an oven for three days at 35, then 40, then 60°C. Embedded nerves were sectioned at 500nm using a Leica Ultramicrotome and a diamond knife (Diatom). Sections were mounted on SuperFrost slides and stained with ρ-phenylenediamine (PPD). Axons were counted using unbiased stereology, described below.

### 2.10 Immunofluorescence and Microscopy

Immediately after sacrifice, eyes were removed and fixed in 4% paraformaldehyde for 30 minutes, then stored in 0.1M PBS. Retinas used for retinal ganglion cell (RGC) counting were dissected from the eye, cryoprotected in 30% sucrose with 0.02% sodium azide for 24 hours, and the vitreous was removed. Whole eyes, after cornea and lens removal, were embedded in OCT (Finetek, Fisher Scientific), sagittally sectioned at 10µm on a cryostat, and sections were placed on SuperFrost slides.

Immunolabeling for RBPMS, the RGC-specific antibody, commenced with 3 freeze-thaw cycles. All tissue was washed three times in 0.1M PBS, incubated in blocking solution (5% donkey serum, 1% Triton X-100 in 0.1M PBS) for one hour, incubated in primary antibody (diluted in blocking solution) for 18-72 hours, washed in PBS, incubated in secondary antibody (diluted in blocking solution), washed in PBS, then coverslipped with either Fluoromount-G or DAPI-Fluoromount-G (Southern Biotech). Primary antibodies and their concentrations are listed in Table 2. Secondary antibodies were from Jackson ImmunoResearch and included Alexa Fluor 488, 594, or 647 conjugated to anti-rabbit, anti-mouse, or anti-goat IgG.

**Table 2.**
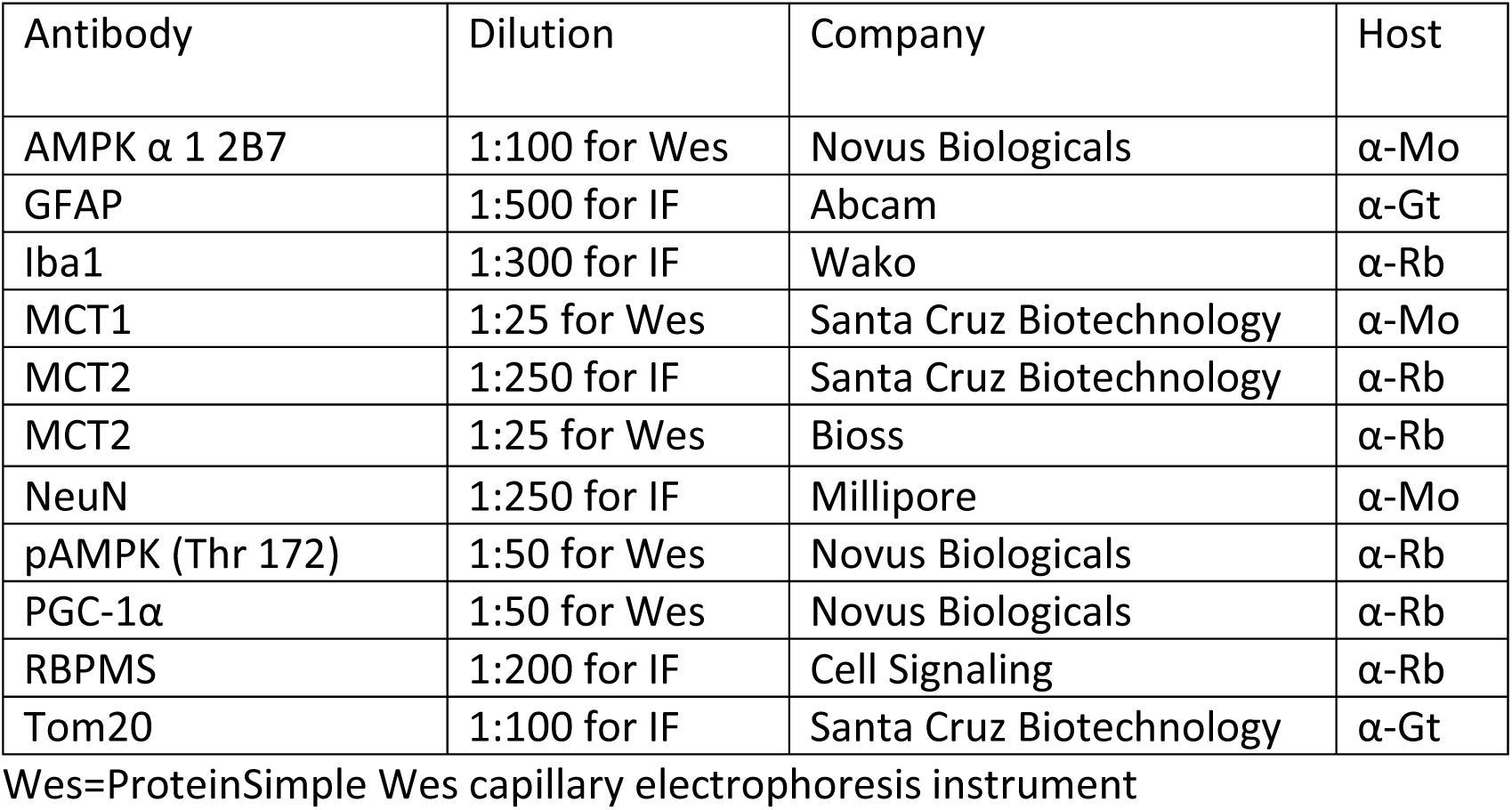
Antibodies Used.

Retinal sections immunolabeled with α-Rb MCT2 primary antibody were incubated with TrueBlack Lipofuscin Autofluorescence Quencher (Biotium) for thirty minutes after blocking and before primary antibody incubation, then washed three times with 0.1M PBS.

For microscopy, a Leica DMi8 confocal microscope (Leica Microsystems, Buffalo Grove, IL, USA) was used. Photomicrographs were captured with consistent exposure and laser intensities among groups. Quantification of Iba1 immunofluorescence was undertaken by measuring fluorescence intensity across microglial cell bodies within regions of interest (ROI) using Fiji-ImageJ. At least 12 ROIs were used per antigen for quantification.

### 2.11 Cell and Axon Quantification

Whole mounted retinas immunolabeled with RBPMS and PPD-stained optic nerve (ON) sections were counted using the optical fractionator approach for unbiased stereology and either a 40x (retina) or 100x (ON) objective (StereoInvestigator software, MBF Bioscience). Between 35 and 40 sampling sites (3%) of each tissue were counted. The coefficient of error (Schmitz-Hof) was kept at 0.05 or below for all stereological counting, ensuring sufficient sampling rate. Iba1 density was calculated by manually counting Iba1-positive cell bodies in the inner retina using a 40x objective, then dividing by the area calculated within Stereoinvestigator. At least 6 sections per group were counted.

### 2.12 Hexokinase Activity

Optic nerves were isolated from sacrificed mice and flash frozen in liquid N_2_. Hexokinase activity was quantified using a hexokinase assay kit (ScienCell), in accordance with the manufacturer’s instructions. Assay results were normalized to the total protein level in each ON sample as quantified with the Pierce BCA protein assay kit (ThermoFisher Scientific).

### 2.13 SDH and COX Histochemistry

Freshly isolated optic nerves (ONs) were flash frozen in liquid N_2_ and held at −80°C until embedding in OCT and sectioning on a cryostat. Control and experimental ONs were sectioned onto the same slide and enzymatically treated simultaneously. All chemicals for histochemistry were obtained from Sigma-Aldrich. ON sections were incubated in 1.5nM nitroblue tetrazolium (N6876), 130mM sodium succinate (S2378), 0.2mM phenazine methosulfate (P9625), and 1mM sodium azide in 0.1M PBS pH=7.0 for 40 min at 37°C for succinate dehydrogenase (SDH) histochemistry. ON sections for cytochrome-c oxidase (COX) were incubated with 1X diaminobenzidine (D4293), 100μM cytochrome c (D2506) and 4 IU/mL catalase (C9322) in 0.1M PBS pH=7.0 for 40min at 37°C. Slides were dehydrated through graded alcohols to xylenes then coverslipped with DPX mounting media.

### 2.14 Protein Analysis

Fresh or flash-frozen ONs were placed in T-PER buffer (Thermo Fisher Scientific) with HALT protease and phosphatase inhibitors (78442, Thermo Fisher Scientific) and homogenized with three 1s pulses at 10% amplitude using a Branson Sonicator. Lysates were centrifuged at 10,000g for 10 min at 4°C; supernatants were collected and used for protein quantification using capillary tube-based immunoassay in the Wes (ProteinSimple). Using the Wes, protein analysis was repeated three times using biological replicates. Wes results are normalized to total protein in the sample, as measured by capillary electrophoresis.

### 2.15 Experimental Design and Statistical Analysis

GraphPad Prism software (version 8.0) was used to generate statistics for quantitative data. Data were analyzed for normality before two-tailed t-test (normally distributed data), or with the non-parametric Mann-Whitney test (non-normally distributed data). One-way ANOVA and Tukey’s multiple comparison post hoc test was used to compare multiple groups within a strain, or across multiple groups within an outcome measure. P<0.05 was considered statistically significant, and data are reported as mean±SD, unless otherwise noted.

## 3. Results

### 3.1 MCT2 Decrease Compromises Visual Function and ATP Production

MCT2 protein is significantly decreased in the glaucomatous retina and ON (Harun-or-Rashid et al., 2018). To evaluate the negative impact on mitochondrial bioenergetics of MCT2 loss, we injected MCT2^fl/+^ mice with an AAV2-cre, generating mice with decreased MCT2 in retinal cells that were infected by the AAV2-cre-GFP. At one and two weeks post-infection, we took fundus photographs to evaluate infection rate and progress. GFP-positive infected cells were observed throughout each injected eye, though the cells were brighter and more distinct at two weeks (Figure 1A). We measured visual evoked potential (VEP) in the mice to determine the functional impact of MCT2^fl/+^. There was a significantly lower response amplitude (N1 trough to P2 peak) in the cre-induced MCT2^fl/+^ mice compared to B6 control (p<0.01; Figure 1B). Representative VEP traces are shown in Figure 1C. The optic nerves from these mice were then evaluated in the Seahorse XF24 Analyzer for oxygen consumption rate (OCR) and extracellular acidification rate (ECAR). MCT2 haploinsufficiency resulted in significantly decreased oxygen consumption for the production of ATP in the cre-induced MCT2^fl/+^ optic nerve (ON) compared to C57Bl/6 (B6) age-matched control, regardless of whether the ONs were in media supplemented with glucose (Glc) or 2-deoxyglucose (2-DG; Figure 1D). The OCR in the B6 ON with 2-DG was no different than when Glc was the substrate, suggesting that there was sufficient lactate or pyruvate stores in the B6 for the ONs to continue ATP production. The cre-induced MCT2^fl/+^ ATP production with either 2-DG or Glc as substrate was no different from the other, but significantly lower than B6 ON, indicating MCT2 haploinsufficiency had a negative overall effect on OCR. In Figure 1E, the extracellular acidification rate (ECAR) was significantly lower in both B6 and MCT2^fl/+^ ON with 2-DG as substrate, reflecting significantly reduced proton release from lactate production.

**Figure 1:**
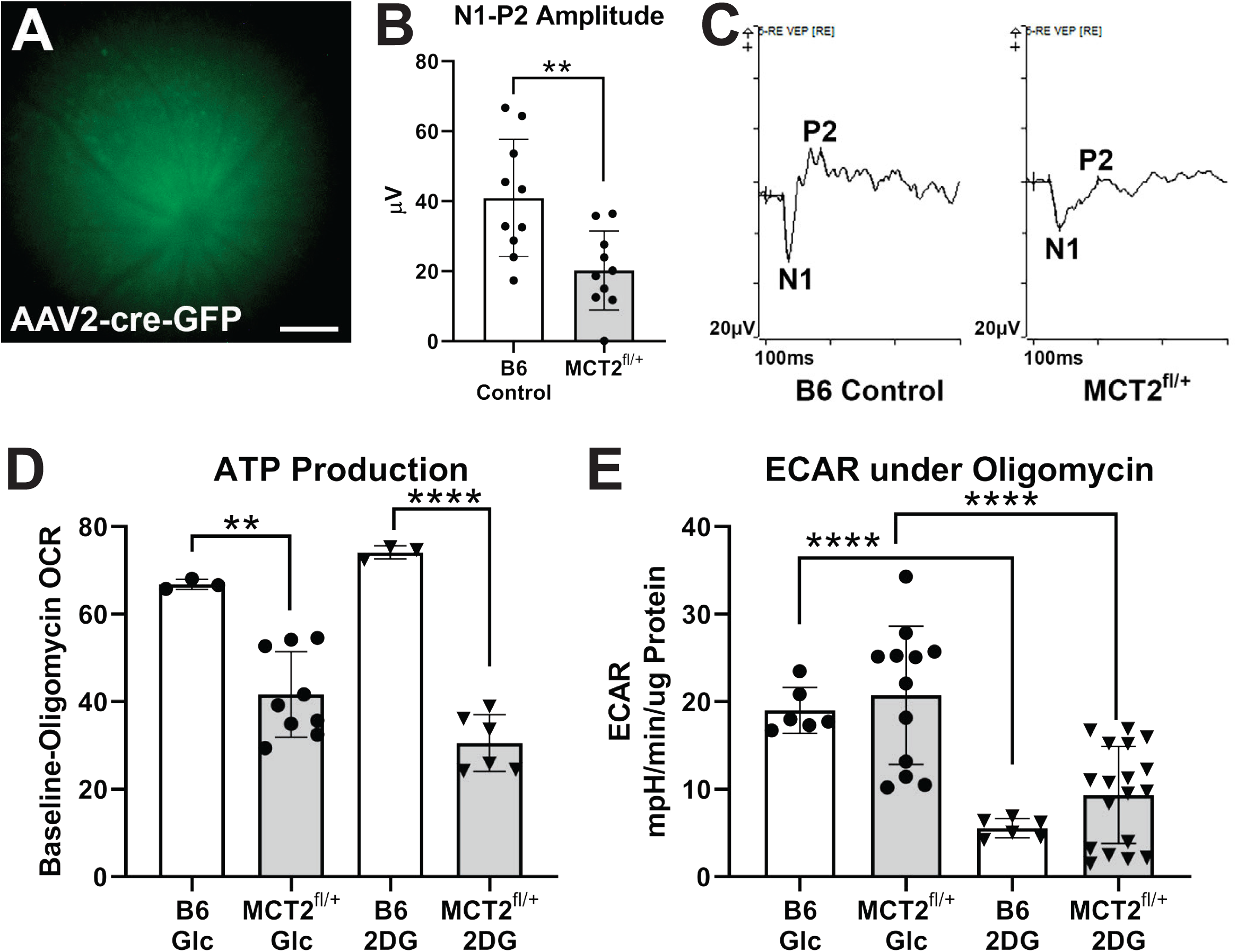
Visual evoked potential (VEP) and optic nerve (ON) metabolism in MCT2^fl/+^ mice. **A**. MCT2^fl/+^ mouse eyes injected with AAV2-cre-GFP were imaged to show expression of GFP in the ganglion cell layer two weeks after infection. Successful cre recombination removes one MCT2 allele from infected cells. Scale bar= 200µm. **B**. VEP response amplitude was significantly reduced in MCT2^fl/+^ mice (p<0.01) compared to age-matched C56Bl/6 (B6) control mice. **C**. Representative VEP traces in B6 control (left) and MCT2^fl/+^ (right) mice. Response amplitude is the µV difference between the N1 trough and the P2 peak. **D**. The difference in baseline versus oligomycin-treatment oxygen consumption rate (OCR) calculated from the ONs of B6 control or MCT2^fl/+^ mice were significantly different for the ONs incubated in glucose (Glc; p<0.1) and 2-deoxyglucose (2-DG; p<0.001). **E**. Extracellular acidification rate (ECAR) was similar for B6 control and MCT2^fl/+^ *within* the Glc or the 2-DG groups, but the 2-DG groups for each were significantly reduced compared to Glc (p<0.0001). Error bars are SD.

### 3.2 MCT2 Overexpression through AAV2:MCT2

Confirming a negative impact on VEP and ATP production with decreased MCT2, we then tested the hypothesis that providing greater access to energy substrates through the upregulation of MCT2 would preserve RGC function. We first immunolabeled control retina to demonstrate localization of MCT2 on RGCs (Figure 2A). The figure panels show retina immunolabeled with antibodies against MCT2, NeuN (neurons), and Vimentin (Müller glia). MCT2 immunolabel is found on RGCs (arrows) and Müller glia endfeet (arrowheads), as well as within the inner plexiform and inner nuclear layers (Figure 2A). In order to upregulate MCT2 in RGCs, we took advantage of the ability of AAV2 to infect neurons after intraocular injection. Figure 2B shows sectioned 12-month-old D2 retina (two months after AAV2:MCT2 injection) with RBPMS-positive RGCs (red), and AAV2:MCT2-positive cells (green from the eGFP reporter). The virus infected both RGCs and neurons in the ganglion cell layer (GCL) and inner nuclear layer (INL) that may be either displaced RGCs or amacrine cells. In some retinas, photoreceptors were observed to be eGFP-positive (data not shown). Figure 2C is the fundus photograph from the AAV2:MCT2 infection in the 10.5-month-old D2 eye two weeks after injection. A further confirmation of the AAV2:MCT2 infection of RGCs is the virus eGFP reporter expression in axons of the optic nerve two months after injection (Figure 2D).

**Figure 2:**
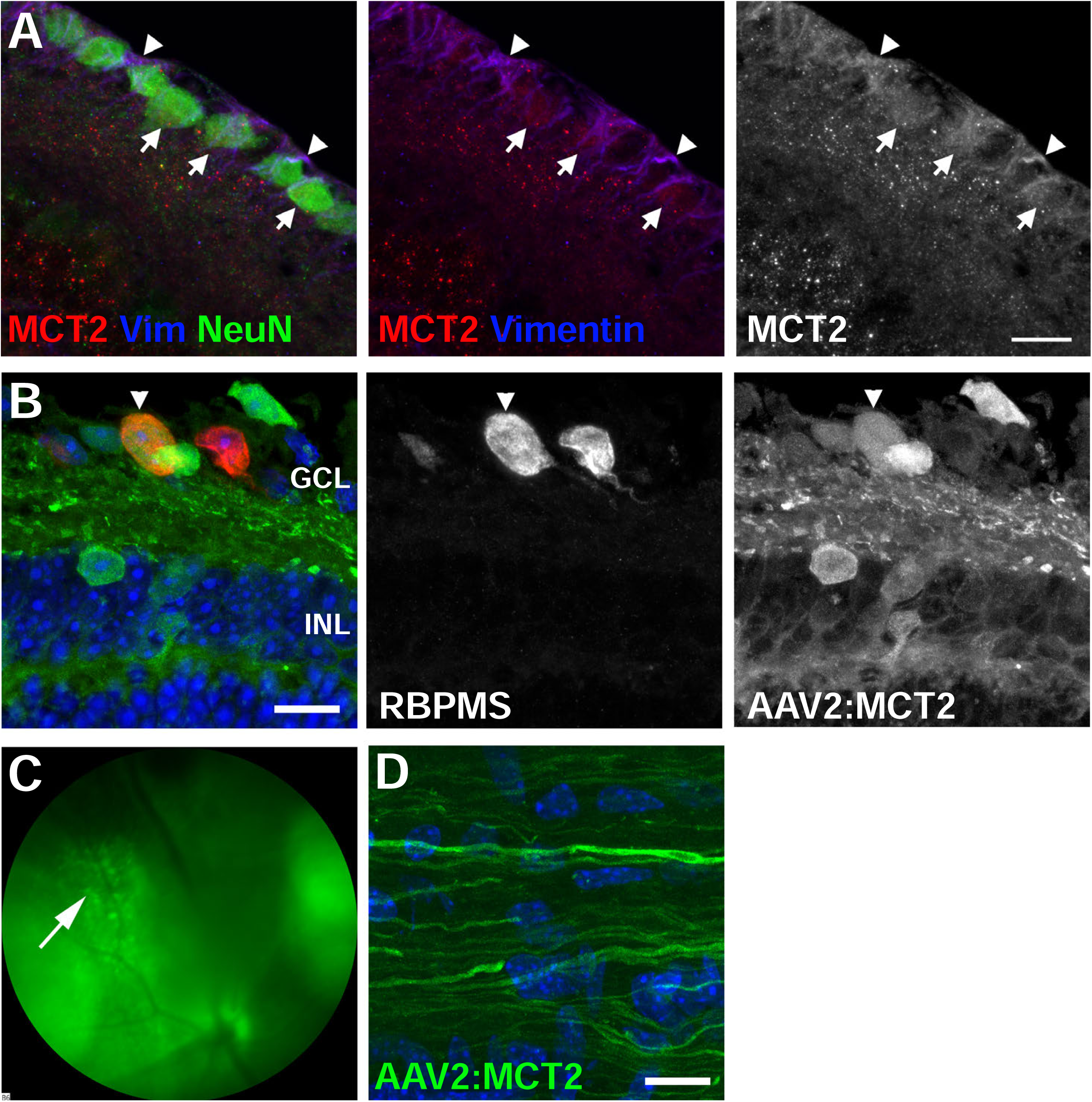
MCT2 protein distribution and AAV2:MCT2 virus infection. **A**. Immunofluorescence of control retina to show MCT2 localization on retinal ganglion cells. The left panel is a merged image of MCT2 (red) with Vimentin (Müller glia, blue) and NeuN (RGCs, green). In the middle panel, NeuN has been removed in order to discern the red puncta along both RGC somas and Müller glia endfeet. The right panel shows MCT2 alone. Scale bar is 15µM. **B**. A cross-section from a 12 month-old D2 retina injected with AAV2:MCT2 two months prior. Arrowheads point to an infected RGC in the GCL that is RBPMS-positive and positive for the eGFP from the AAV2:MCT2 virus. INL=inner nuclear layer; OPL=outer plexiform layer. Scale bar=15µm. **C**. Fundus photograph of a 10.5 month-old DBA/2J (D2) eye injected with AAV2:MCT2, two weeks after injection (left). The arrow points to GFP-positive cells in the ganglion cell layer (GCL). **D**. Section of a 12 month-old D2 optic nerve from an eye injected with AAV2:MCT2. The eGFP-positive axons confirm the infection of RGCs with the virus and the widespread transport of the eGFP. Scale bar=20µm.

### 3.3 MCT2 Overexpression Protects RGCs from Degeneration

Mean D2 mouse IOP for all groups (untreated, AAV2 with or without ketogenic diet) at 10 months of age and prior to intervention was 21.7±0.75; at 12 months of age, prior to sacrifice, mean IOP was 22.5±0.9. There was no apparent impact on IOP of the AAV2:MCT2 injection. Treated D2 mice were injected with AAV2:MCT2 at 10 months of age. As previous data in our laboratory demonstrated that placing D2 mice on a ketogenic diet for two months resulted in upregulation of MCT2, we sought to probe whether diet would be necessary for the function and maintenance of the MCT2 after viral-mediated overexpression. Therefore, we included an AAV2:MCT2 injected group of D2 mice also on the ketogenic diet. The AAV2:MCT2 injected eyes, regardless of diet, had significantly greater RGC density than the 12-month-old age-matched untreated D2 retina (Figure 3A, p<0.0001). Images of retinas immunolabeled with RBPMS for RGCs showed RGC loss in the untreated D2 retina (Figure 3B). Cross-sections of optic nerves stained with ρ-phenylenediamine (Figure 3C) show loss of axons in the untreated D2 and in the AAV2:MCT2 D2 mice on the ketogenic diet (KD). The AAV2:MCT2-injected D2 mice on a normal diet had significantly greater axon number in the optic nerve (ON) than untreated D2 (Figure 3D, p<0.0001). However, the 12-month-old D2 mice on the ketogenic diet showed significantly lower axon number than D2 mice on the control diet (p<0.05), despite a few animals with axon number in the normal range.

**Figure 3:**
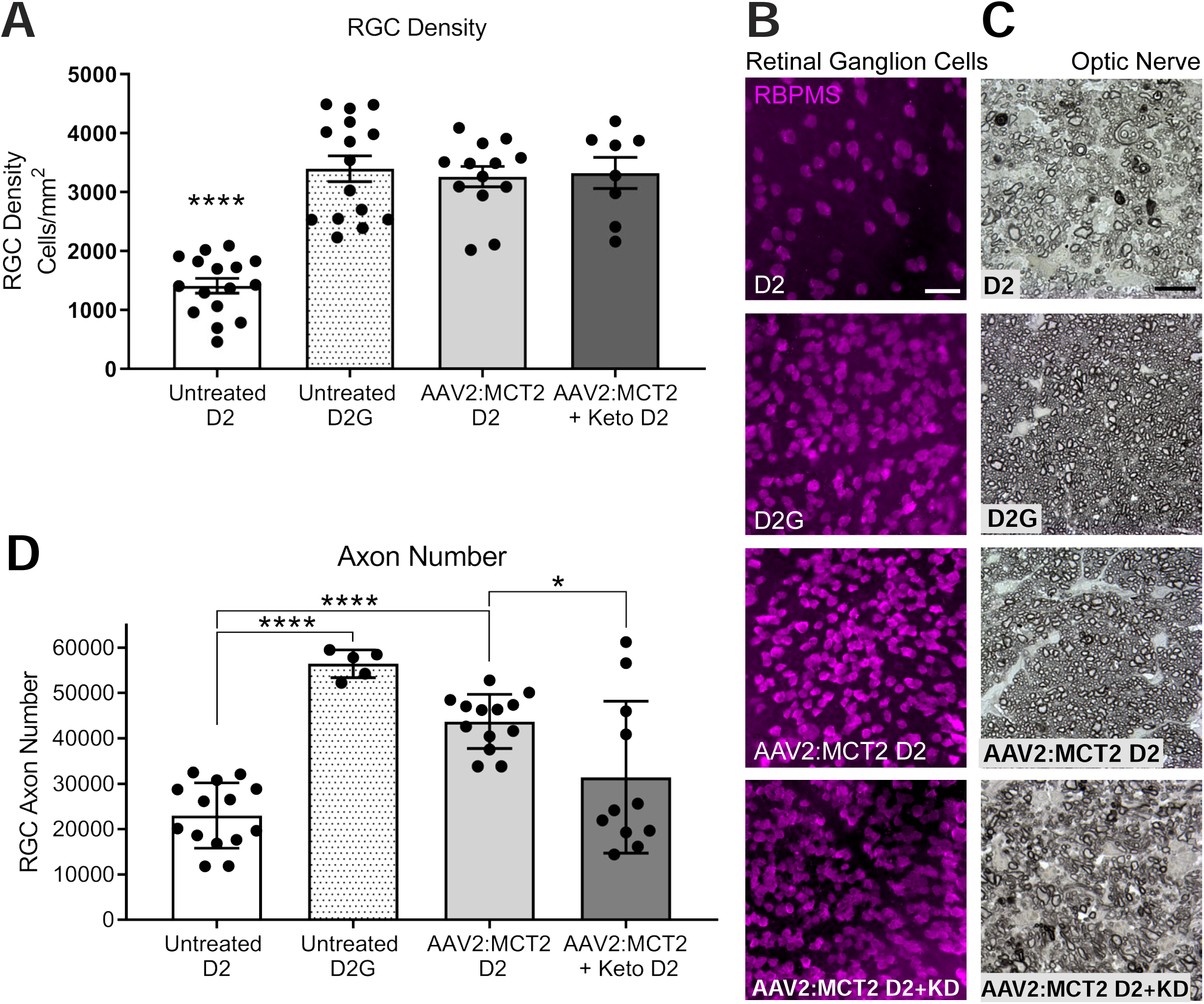
Retinal ganglion cell soma and axon quantification and integrity. **A**. Retinal ganglion cell density (cells/mm^2^) in 12 month-old D2 retina, age-matched control strain DBA/2-*Gpnmb*^+^ (D2G) retina, 12 month-old D2 retina injected with AAV2:MCT2, and 12 month-old D2 retina injected with AAV2:MCT2 and placed on a ketogenic diet for 8 weeks. D2G and both virus injected groups have significantly greater RGC density than 12 month-old D2 (p<0.0001). **B**. Wholemount retina immunolabeled for RBPMS, an RGC-specific marker, from untreated D2 and D2G retina, and D2 mouse retina injected with AAV2:MCT2, with or without the ketogenic diet (KD). Images were taken 1mm from the optic nerve head. Scale bar=25µm. **C**. Cross-sections of ρ-phenylenediamine labeled optic nerve from untreated D2 and D2G, then D2 mice injected with AAV2:MCT2, with or without the ketogenic diet (KD). Scale bar=10µm. **D**. Optic nerve (ON) axon counts from D2, D2G, and D2 injected with AAV2:MCT2, with or without the ketogenic diet. D2 axon number is significantly lower than D2G and AAV2:MCT2 D2 (p<0.0001). AAV2:MCT2 D2 mice on the ketogenic diet had significantly lower ON axon number than the AAV2:MCT2 D2 mice on normal rodent chow (p<0.05). Error bars are SD.

### 3.4 MCT2 Overexpression Reduced Increased Mitochondrial Activity

To confirm upregulation of MCT2 protein through viral-mediated overexpression, we used capillary electrophoresis to quantify MCT2 in the ON, finding MCT2 levels that matched the D2G mouse ON and were significantly greater than the untreated 12-month-old D2 ON (Figure 4A, p<0.0001). We evaluated whether MCT2 upregulation would have a compensatory effect on MCT1 expression, so we quantified MCT1 in the ON. MCT1 protein is significantly higher in D2G and D2 mice injected with AAV2:MCT2 than untreated D2 mice (Figure 4B, p<0.05). The ratio of phosphorylated AMP kinase (pAMPK) to AMP kinase increases when ATP levels are low. The D2 mice have significantly higher pAMPK/AMPK ratio in the ON than either the D2G control or the D2 injected with AAV2:MCT2 (Figure 4C, p<0.001). PGC-1α (PPARγ coactivator 1) stimulates mitochondrial biogenesis and plays a central role in cellular metabolism, so we anticipated it may be altered by MCT2 overexpression. PGC-1α protein in ON from D2 mice infected with AAV2:MCT2 is significantly increased compared to both untreated D2 and D2G mice (Figure 4D, p<0.01). Upregulation of MCT2 may impact the balance of energy derived from glycolysis versus oxidative phosphorylation, so we evaluated hexokinase activity, the first enzyme in the glycolytic pathway and a key regulator of glycolytic flux (Tanner et al., 2018). Hexokinase activity is upregulated in D2 mice injected with AAV2:MCT2 compared to untreated D2 mice (Figure 4E, p<0.05). Succinate dehydrogenase (SDH) activity, contributing to both the TCA cycle and the electron transport chain, can be a proxy for cellular respiration rate. We observed that SDH activity in ON cross-sections subjected to SDH histochemistry is significantly higher in D2G and D2 mice injected with AAV2:MCT2 than untreated D2 (Figure 4F, p<0.0001).

**Figure 4:**
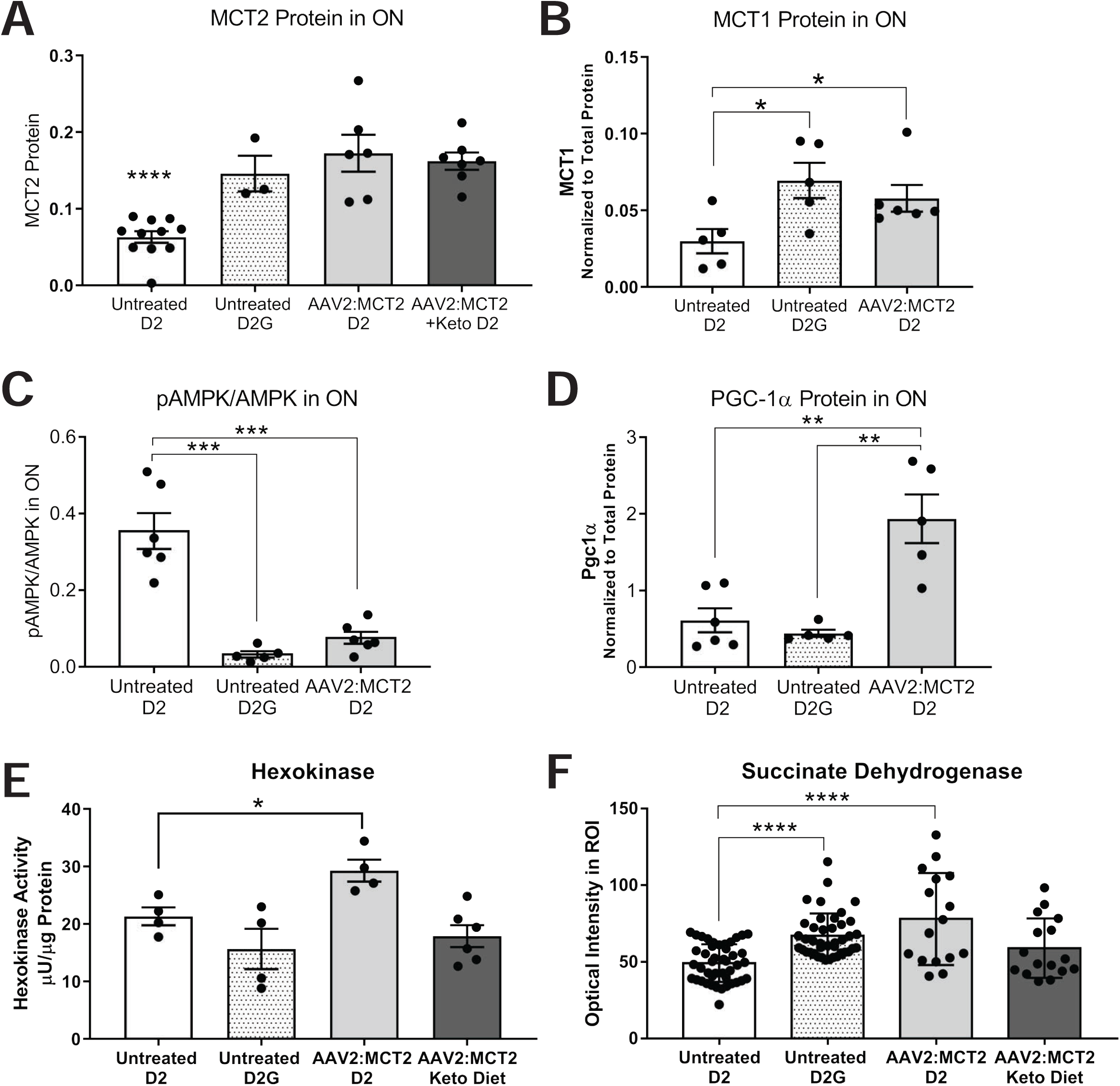
Protein and metabolic changes in optic nerve (ON) from D2 mice injected with AAV2:MCT2. **A**. MCT2 protein in ON from D2, D2G mice, and D2 mice injected with AAV2:MCT2, with or without the ketogenic diet. Untreated D2 MCT2 protein is significantly lower (p<0.0001) than each of the other experimental groups depicted. **B**. MCT1 expression is higher in both untreated D2G and AAV2:MCT2 injected D2 than in untreated D2 optic nerve (p<0.05). **C**. PGC-1α is upregulated in D2 mice injected with AAV2:MCT2 compared to both untreated D2 and D2G mice (p<0.01). **D**. pAMPK/AMPK ratio is significantly higher in untreated D2 than D2G and AAV2:MCT2 injected D2 mice (p<0.001). **E**. Hexokinase activity is significantly increased in AAV2:MCT2 injected D2 mice compared to untreated D2 (p<0.05). **F**. Succinate dehydrogenase activity is significantly decreased in untreated D2 mouse ON compared to both untreated D2G and AAV2:MCT2 treated D2 mice on standard chow (control diet), p<0.0001. Error bars are SD.

### 3.5 Microglial Activation Increased with AAV2:MCT2 Injection

Virus injection can induce an inflammatory response, so we evaluated microglia in the experimental retinas. Iba1-positive microglia are observed in all retinas, primarily in the outer plexiform, inner plexiform, and ganglion cell layers (Figure 5A-D). In the untreated 12 month-old D2 retina (Figure 5A), the Iba1-positive microglia are intensely labeled and less ramified than the D2G control retinas (Figure 5B). Numbers of microglia and their Iba1 labeling intensity increase in both groups injected with AAV2:MCT2 (D2 mice ± ketogenic diet), Figure 5C and D. Both AAV2:MCT2 injected D2 ± ketogenic diet show significantly higher microglial density than untreated D2 and D2G mouse retina (Figure 5E, p<0.0001). Iba1 fluorescence intensity, correlated to the level of microglial activation (Bosco et al., 2008), is highest in the AAV2:MCT2 injected D2 retina on the control diet. Untreated D2 retina has significantly higher Iba1 fluorescence intensity compared to untreated D2G mice (Figure 5F, p<0.001). D2 mice injected with AAV2:MCT2 ± the ketogenic diet have higher Iba1 activation than untreated D2 mice (Figure 5F, p<0.05 and p<0.001).

**Figure 5:**
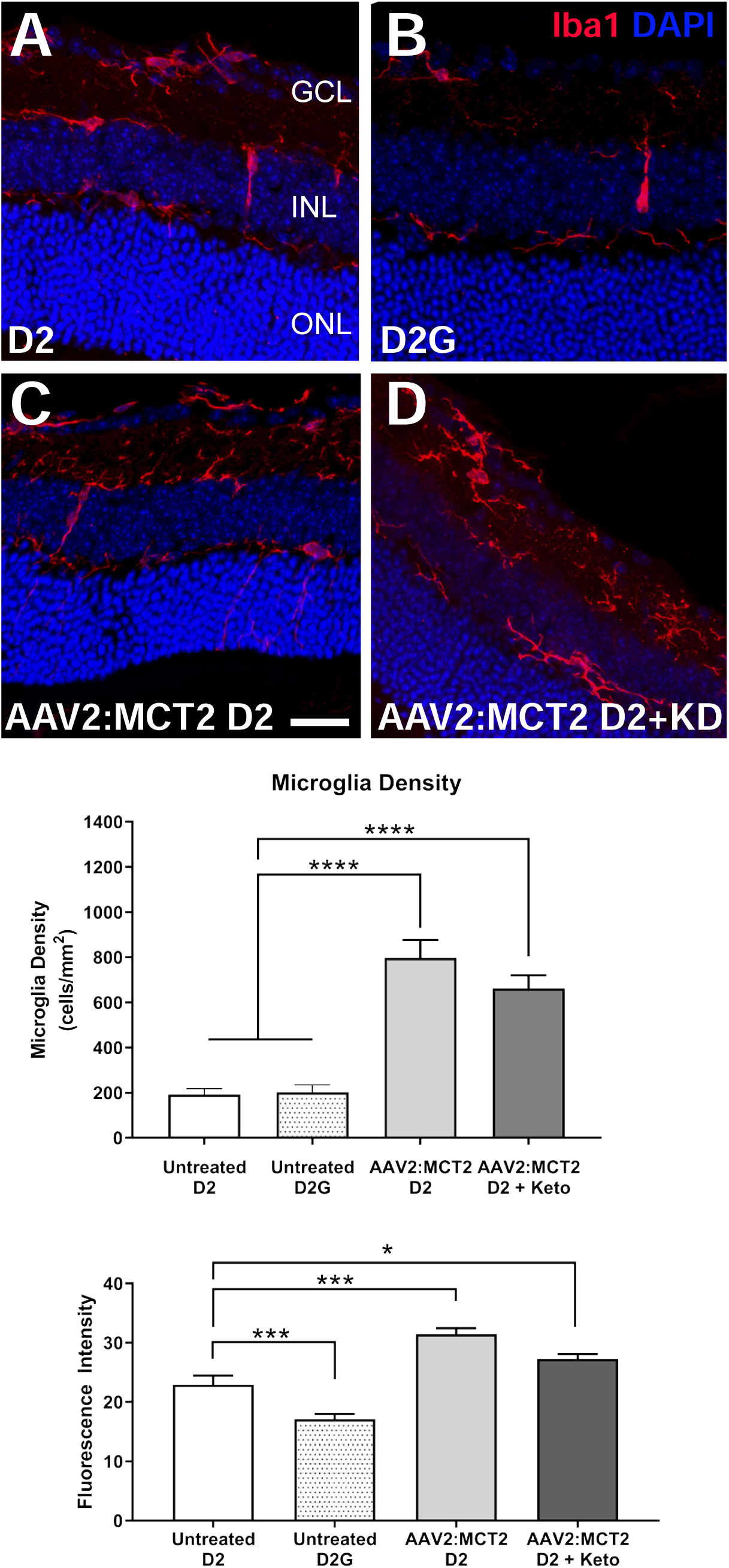
Iba1 fluorescence and microglia density in retinas from D2 mice injected with AAV2:MCT2. **A-D**. Photomicrographs showing the expression of Iba1 (red) in microglia by immunolabeling in retinas from D2 (**A**), D2G (**B**), D2 injected with AAV2:MCT2 (**C**), and D2 injected with AAV2:MCT2 mice on the ketogenic diet (KD) (**D**). Scale bar = 35µm. DAPI nuclear stain in blue. **E**. D2 mice injected with AAV2:MCT2 with or without ketogenic diet showed significantly higher microglia density than untreated D2 and D2G mice (p<0.0001). **F**. Untreated D2 mice had significantly higher Iba1 fluorescence than untreated D2G mice (p<0.001), but significantly lower Iba1 fluorescence than either AAV2:MCT2 injected D2 mice (p<0.001) or AAV2:MCT2 injected D2 mice on the ketogenic diet (p<0.05). Error bars are SEM.

### 3.6 MCT2 Overexpression Rescued RGCs after Ocular Hypertension

To further explore the effects of increasing MCT2 expression in the context of glaucoma, we undertook an analysis of mice injected with AAV2:MCT2 after being subjected to ocular hypertension. Figure 6A shows flatmount retina from a Thy1-tdTomato mouse injected with AAV2:MCT2. There is widespread colocalization of Thy1-positive RGCs and the eGFP from the virus reporter. Figure 6B shows the intraocular pressure (IOP) profiles for the OHT groups: OHT with AAV2:MCT2 injection, OHT with control virus (AAV2:eGFP) injection, and OHT alone. All OHT groups underwent significant increases in IOP (p<0.001) compared to baseline values. Mice were held until each achieved 200mmHg-Days of IOP increase, hence the range in weeks of IOP measurement. There were no statistical differences across OHT groups for peak IOP or IOP integral. RGC density is significantly higher in OHT mice injected with AAV2:MCT2 and control mice than in OHT mice (Figure 6C, p<0.01). ON axon number is also significantly higher in OHT mice injected with AAV2:MCT2 and control mice than in OHT mice (Figure 6D, p<0.001, and p<0.01, respectively). OHT mice injected with AAV2:MCT2 have significantly higher MCT2 protein in retina than both control and OHT retina (Figure 6E, p<0.001 and p<0.0001). In ON, there is a significant effect of group on MCT2 protein, though no individual statistical differences were observed (ANOVA, p<0.05, Figure 6F).

**Figure 6:**
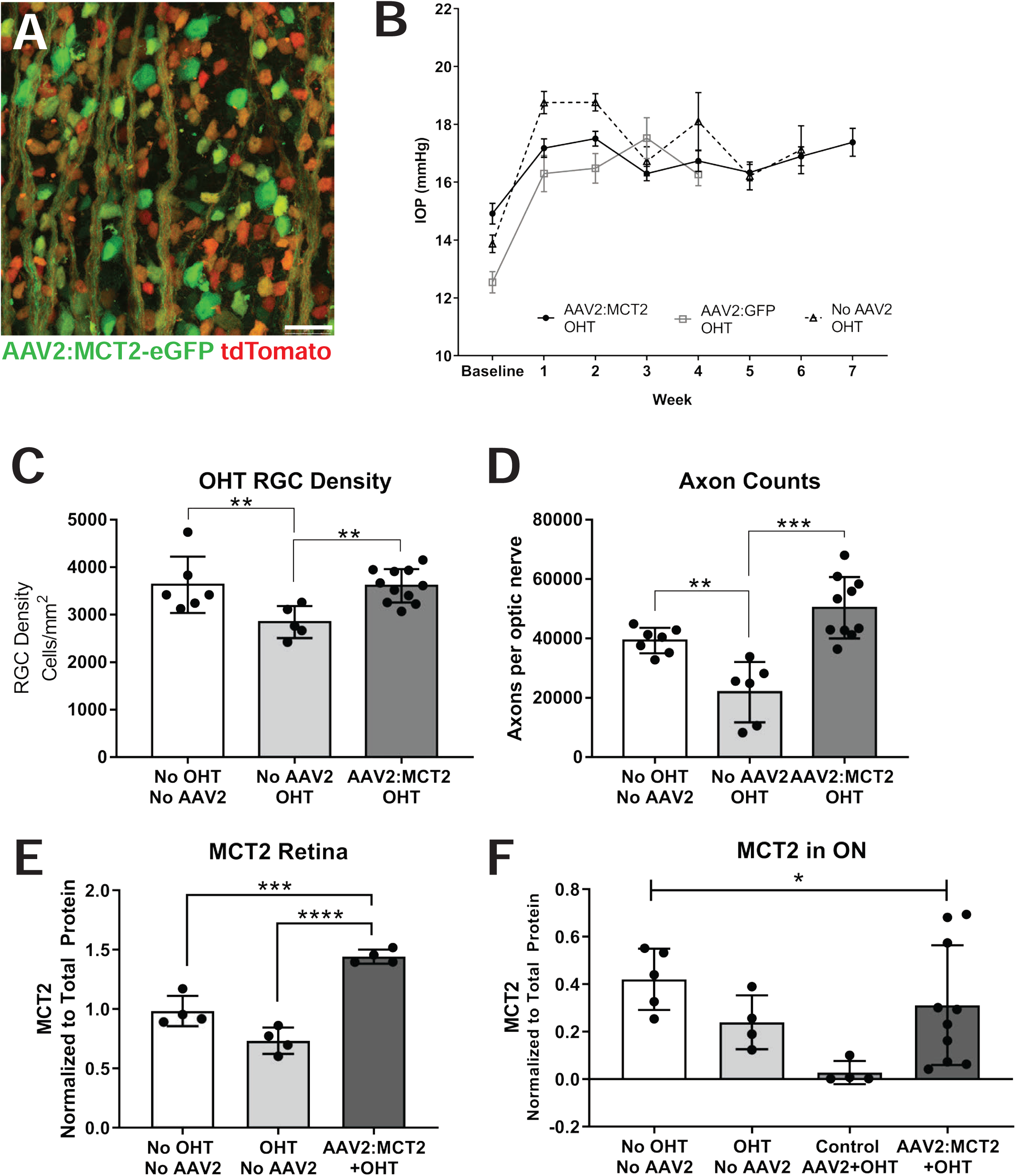
Retinal ganglion cell (RGC) and optic nerve (ON) axon integrity, and MCT2 protein levels in mice after ocular hypertension (OHT). **A**. Photomicrograph of AAV2:MCT2 infection in flatmount retina without OHT; AAV2:MCT2 virus infection reporter is eGFP (green), and RGC-specific label RBPMS (red). Scale bar = 50µm. **B**. Intraocular pressure (IOP) elevation in the three OHT experimental groups. Each OHT group IOP was significantly increased over baseline, p<0.0001. Error bars are SEM. **C**. RGC density is significantly increased by AAV2:MCT2 injection after OHT compared to OHT alone (p<0.01). OHT alone resulted in significant RGC loss compared to control (no OHT and no AAV2; p<0.01). **D**. Axon number in the ON of OHT mice is significantly decreased compared to control (No OHT, p<0.001), then rescued by injection with AAV2:MCT2 after OHT (p<0.001). **E**. MCT2 protein in retina is significantly higher in the OHT group injected with AAV2:MCT2 compared to both control (No OHT; p<0.001) and OHT alone (p<0.0001). **F**. MCT2 protein in ON is significantly different across groups by one-way ANOVA (p<0.05). Error bars are SD.

### 3.7 MCT2 Overexpression Reduced Metabolic Stress after OHT

AAV2:MCT2 infection changes the metabolic activity in OHT mice. OHT mice injected with AAV2:MCT2 have a significantly lower pAMPK/AMPK ratio in ON than control mice, and OHT mice injected with AAV2:eGFP (Figure 7A; p<0.01, and p<0.05 respectively). PGC-1α protein in ON shows no change across control and experimental groups (Figure 7B). In contrast to the D2 experiments which showed significantly greater hexokinase activity with AAV2:MCT2 infection, hexokinase activity is significantly lower in OHT mice injected with AAV2:MCT2 compared to OHT alone (Figure 7C, p<0.05). Mitochondrial size is not different between OHT mice with either AAV2:eGFP or AAV2:MCT2 infection (Figure 7D).

**Figure 7:**
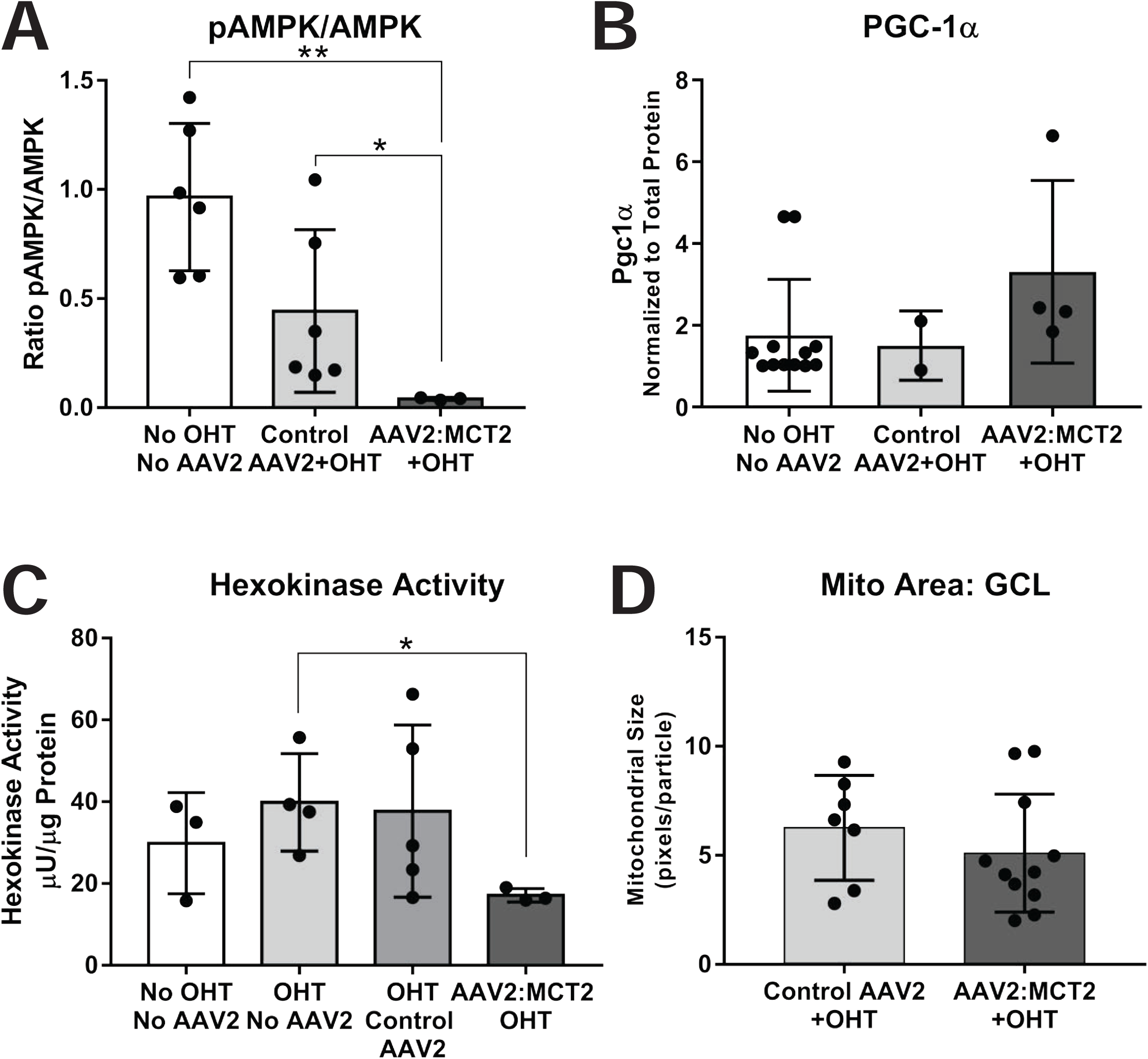
Metabolic changes in ON from OHT mice injected with AAV2:MCT2. **A**. The ratio of phosphorylated AMP kinase (pAMPK) to AMPK is significantly lower in OHT mice injected with AAV2:MCT2 compared to control mice (no OHT or virus) and OHT mice injected with AAV2:GFP (**p<0.01, *p<0.05 respectively). **B**. PGC-1α protein in ON did not differ statistically across groups. **C**. Hexokinase activity is significantly decreased in OHT mice injected with AAV2:MCT2 compared to OHT mice alone (*p<0.05). **D**. Mitochondria size in the ganglion cell layer (GCL) did not differ across virus groups. Error bars are SD.

### 3.8 MCT2 Overexpression Increased Mitochondrial Function after OHT

COX and SDH activity are significantly increased in OHT mice injected with AAV2:MCT2 compared to OHT alone. Lower COX activity in OHT control ON compared to the AAV2:MCT2 OHT group is evident in the high magnification insets of Figure 8A (compare a’ to b’). SDH is also less active in OHT mice (Figure 8C, c’) than in OHT mice injected with AAV2:MCT2 (Figure 8D, d’). OHT mice injected with AAV2:MCT2 have significantly lower SDH activity than control mice (Figure 8E, p<0.001), but significantly higher SDH activity than OHT alone and OHT mice injected with AAV2:eGFP (Figure 8E; p<0.001, p<0.05 respectively). OHT mice injected with AAV2:MCT2 also have significantly higher COX activity than OHT mice (Figure 8F, p<0.0001).

**Figure 8:**
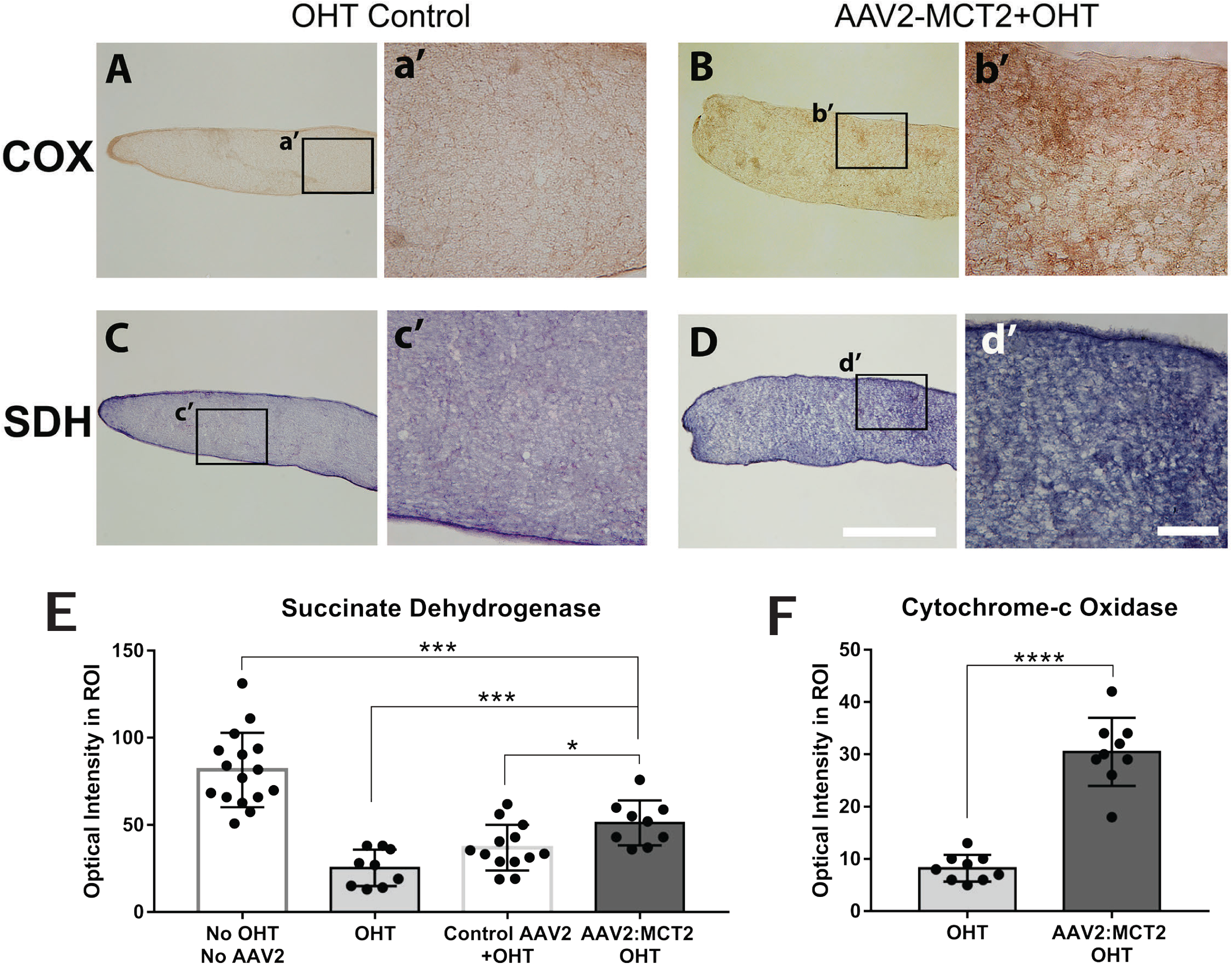
Cytochrome-c oxidase (COX) and succinate dehydrogenase (SDH) activity in OHT optic nerves. **A-B**. COX histochemistry in OHT mouse ON (**A**), and OHT with AAV2:MCT2 injection (**B**). Higher magnification insets of OHT control (**a’**) and AAV2:MCT2 + OHT (**b’**) show greater DAB intensity after OHT with AAV2:MCT2. Quantification of histochemistry is shown in F. **C-D**. SDH histochemistry in OHT mouse ON (**C**) and OHT with AAV2:MCT2 injection (**D**). Higher magnification insets of OHT control (**c’**) and AAV2:MCT2 + OHT (**d’**) indicate greater nitroblue tetrazolium intensity after OHT with AAV2:MCT2. **E**. SDH activity is significantly higher in AAV2:MCT2 with OHT ON compared to both OHT alone, and OHT ON injected with control AAV2 (AAV2:GFP) (p<0.001 and p<0.05, respectively). SDH activity is significantly lower in OHT animals injected with AAV2:MCT2 than in control mice (no OHT or AAV2). **F**. COX activity is significantly higher in OHT mouse ON injected with AAV2:MCT2 compared to OHT alone (p<0.0001). Error bars are SD.

### 3.9 MCT2 Overexpression Protected Inner Retinal Function after OHT

Microglia density does not differ between OHT mice injected with AAV2:eGFP or AAV2:MCT2 (Figure 9A). However, there is a significant effect of virus injection lowering Iba1 fluorescence intensity across the OHT groups (Figure 9B, p<0.0001). OHT mice injected with AAV2:MCT2 had significantly higher pattern electroretinogram P1 peak amplitude than OHT mice (Figure 9C, p<0.05). N2 amplitude shows no difference across the control and experimental groups (Figure 9D). Representative PERG traces for each analyzed group are shown in Figure 9E.

**Figure 9:**
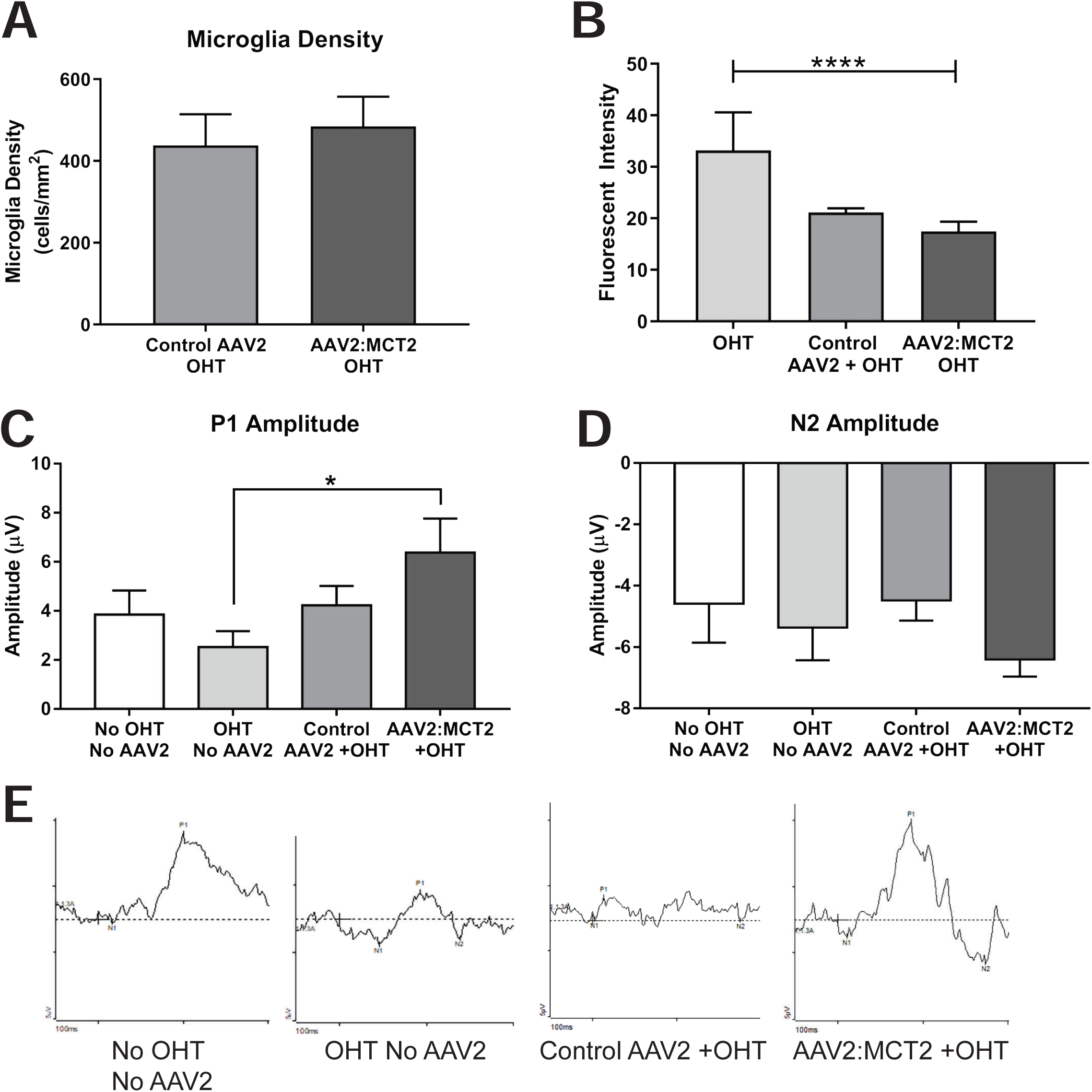
Retinal microglia and synapse analysis, and pattern electroretinogram in OHT mice. **A**. Microglia density did not differ between OHT eyes injected with control virus (AAV2:eGFP) or AAV2:MCT2. **B**. Intensity of Iba1 immunolabeling was significantly different across groups, with virus injection having a lowering effect on fluorescence intensity of Iba1 (ANOVA, p<0.0001). **C**. Pattern electroretinogram shows significantly higher P1 amplitude in OHT animals injected with AAV2:MCT2 than in OHT alone (p<0.05). **D**. N2 amplitude did not differ across experimental groups. Error bars are SD. **E**. Example PERG traces for each of the groups represented in **C** and **D**.

## 4. Discussion

Reduction of MCT2 in the retina through cre-mediated elimination of flox’d MCT2 alleles resulted in compromised ATP production and visual evoked potential. These findings corroborate our observations of the significant, negative impact of MCT2 decrease in mouse models of glaucoma (Harun-or-Rashid et al., 2018), and motivated our strategy to increase metabolic substrate availability in the visual system through overexpression of MCT2. Using AAV2 to increase MCT2 expression in the retina and ON of D2 mice and those with OHT resulted in rescue of RGC soma and axon number, restoration of energy homeostasis, and increased mitochondrial activity.

The kinetics of MCT2 indicates the transporter has a high affinity for monocarboxylates (Bröer et al., 1999), a quality associated with a cell type that imports, rather than exports, monocarboxylates. RGCs and Müller glia are likely importers of monocarboxylates, the latter because their pyruvate kinase deficiency necessitates carbon sources others than glucose to fuel their oxidative phosphorylation (Lindsay et al., 2014). Gerhart et al, concluded that MCT2 was only associated with glial cells despite showing immunolabeling that included RGCs (Gerhart et al., 1999). We have demonstrated MCT2 expression on RGCs and their axons (Harun-or-Rashid et al., 2018) and adopted an AAV2-mediated infection strategy that was designed to overexpress MCT2 in RGCs. We observed no AAV2-associated GFP reporter expression in cells other than neurons in the retina.

Since MCT2 transports ketone bodies in addition to lactate and pyruvate, we asked ourselves whether greater availability of ketone bodies in the bloodstream would be necessary for the potential improvement in energy homeostasis in the AAV2:MCT2 injected mice. Significant protection of RGC somata was observed in D2 mice on the ketogenic diet injected with AAV2:MCT2; however, we observed a negative impact on axon number of the ketogenic diet on the D2 mice with AAV2:MCT2 infection. There were a few optic nerves with normal numbers of RGC axons, but the majority showed no difference in axon number from untreated D2 mice. This was surprising as it suggests the viral overexpression counteracted the D2 axon protection observed with the ketogenic diet in past publications (Harun-or-Rashid et al., 2018). One potential explanation was the significant increase in Iba1+ cell number and fluorescence intensity, an indication of microglial activation, that was observed in the mice injected with AAV2:MCT2. A prominent effect of the ketogenic diet is its inhibition of NF-κB signaling and the NLRP3 inflammasome through HCAR1, a ketone body receptor expressed on microglia, among other cells (Harun-or-Rashid and Inman, 2018). It is likely that the virus-mediated inflammatory response overcame the anti-inflammatory effect of the ketogenic diet, resulting in greater axon loss. One important difference between our past ketogenic study and this one was the age of the D2 mice; they were 11 months of age at analysis in the prior study, but 12 months of age in this study, so likely to have a more progressed glaucoma pathology. Given the magnitude of the RGC protection in older mice here, the MCT2 overexpression may be of greater benefit than the ketogenic diet to RGC survival and function. Ultimately, we were able to determine that the virus-mediated MCT2 expression did not require the diet to be effective.

It was our expectation that we would observe a decline in hexokinase activity with AAV2:MCT2 injection because we hypothesized that increased MCT2 would lead to greater monocarboxylate import, increased mitochondrial function and lowered glycolysis. The OHT mice injected with AAV2:MCT2 had significantly lower hexokinase activity than OHT alone. Since control virus injected OHT eyes had similar hexokinase activity as the OHT alone, it was evident that the addition of MCT2 was associated with the decline in hexokinase activity. The AAV2:MCT2 OHT ON achieved energy homeostasis (low pAMPK/AMPK ratio), and the SDH and COX activities were significantly greater than OHT alone, indicating that mitochondrial function was indeed higher in the AAV2:MCT2 OHT ON. However, hexokinase activity was significantly upregulated in the AAV2:MCT2 D2 ON compared to untreated D2. For the D2, it may be the case that the improvement in energy homeostasis (evident with the decline in pAMPK/AMPK ratio) enabled improved function for all modes of ATP generation, glycolysis and oxidative phosphorylation. Another possibility is that the D2 ON, relying heavily on glycolysis as we have demonstrated in prior studies (Jassim et al., 2019), would not have experienced an increase in monocarboxylate import through MCT2 sufficient enough to displace the energy that could be obtained through glycolysis. Important to note is that the observed changes in hexokinase activity are relative; the hexokinase activity in the D2 ON with AAV2:MCT2, despite being significantly higher than untreated D2, was still lower than the control ON in the OHT experiment.

PGC-1α was significantly upregulated in ON after AAV2:MCT2 injection in retina, for both models of glaucoma. PGC1α can contribute in significant ways to cellular protection in glaucoma (Guo et al., 2014; Harun-or-Rashid et al., 2018) by suppressing reactive oxygen species (ROS) generation through expression of ROS scavenging enzymes such as SOD2 (St-Pierre et al., 2006). PGC1α also coordinates mitochondrial biogenesis by coactivating TFAM (mitochondrial transcription factor A), a direct regulator of mtDNA transcription (Wu et al., 1999). Relatedly, we observed significant increases in succinate dehydrogenase (SDH) activity in the D2 with AAV2:MCT2 injection. SDH is part of the TCA cycle and the mitochondrial electron transport chain; its activity is a proxy for mitochondrial function. The high SDH and cytochrome c oxidase activities in ONs from AAV2:MCT2 injected mice indicate MCT2 overexpression allowed for import of various energy substrates that could be utilized by mitochondria for TCA cycle and electron transport chain function.

One drawback of viral vectors is the possibility of engendering an inflammatory response. Microglial density increased in D2 retina with AAV2:MCT2 infection, regardless of diet. There was no change in microglial density in OHT retina with or without AAV2:MCT2 infection. This difference between D2 and OHT microglia density was not unexpected for the D2, given the significant inflammatory response observed in the D2 retina (Bosco et al., 2011, 2008). Toll-like receptors mediate the initial immune response to AAV (Zhu et al., 2009), including NF-κB activation with downstream cytokine production (Rogers et al., 2011). Cytokines such as TNFα, IL-6, IFNα/β, and MCP-2 released from AAV capsid-ingesting antigen presenting cells (Rogers et al., 2011) have been observed in glaucomatous retina and ON (Chidlow et al., 2012; Tezel et al., 2001; Wilson et al., 2015). Despite the limited data on antigen presenting cells in retina, the potential for additional cytokine release raises the possibility that AAV injection would exacerbate immune response there. However, the absence of microglial density change and the significant decline in microglial Iba1 intensity with AAV2:MCT2 after OHT suggest the virus was not as directly immunogenic as it appeared to be with the D2 mouse. As mentioned above, the virus infection was likely to have reversed the anti-inflammatory effect of the ketogenic diet.

MCT inhibition using α-cyano-4-hydroxycinnamate resulted in a reduction of the flash ERG b-wave and associated oscillatory potentials (B. V. Bui et al., 2004), suggesting that glutamatergic neurotransmission from photoreceptors to bipolar cells was negatively impacted by inhibition of monocarboxylate transport. The implication is that glutamate would have been shunted from the glutamate-glutamine cycle to contributing to the TCA cycle. In this study, pattern electroretinogram showed a P1 amplitude that was significantly higher in the AAV2:MCT2 mouse with OHT compared to OHT alone. There was a trend toward greater N2 amplitude with AAV2:MCT2 infection, but it was not statistically different from control or OHT alone. The significance of the P1 amplitude is unknown since the source of the individual components of the PERG waveform has not been established (Porciatti, 2007).

## 5. Conclusion

Facilitating the transport of monocarboxylates in the retina during ocular hypertension, either through short-term (4 weeks) or long-term (6 months) IOP increase, maintained RGC number, improved energy homeostasis, and increased mitochondrial function in the retina and ON. These findings indicate metabolic intervention to support visual function is a viable approach for the treatment of glaucoma.

## Acknowledgements

The expert technical assistance of Amelia McMullen and Josephine Lepp are gratefully acknowledged.

